# Optogenetic control of apical constriction induces synthetic morphogenesis in mammalian tissues

**DOI:** 10.1101/2021.04.20.440475

**Authors:** Guillermo Martínez-Ara, Núria Taberner, Mami Takayama, Elissavet Sandaltzopoulou, Casandra E. Villava, Nozomu Takata, Mototsugu Eiraku, Miki Ebisuya

**Author notes:** Correspondence to M Ebisuya. These authors contributed equally to this work (Guillermo Martínez-Ara, Núria Taberner, Mami Takayama).

## Abstract

During embryonic development, cellular forces synchronize in space and time to generate functional tissue shapes. Apical constriction is one of these force-generating processes, and it is necessary to modulate epithelial curvature in fundamental morphogenetic events, such as neural tube folding. The emerging field of synthetic developmental biology proposes bottom-up approaches to examine the contribution of each cellular process to complex morphogenesis. However, the shortage of tools to manipulate three-dimensional (3D) shapes of mammalian tissues currently hinders the progress of the field. Here we report the development of “OptoShroom3”, a new optogenetic tool that achieves fast spatiotemporal control of apical constriction in mammalian epithelia. Activation of OptoShroom3 through illumination of individual cells in an epithelial cell sheet reduced their apical surface while illumination of groups of cells caused deformation in the adjacent regions. By using OptoShroom3, we further manipulated 3D tissue shapes. Light-induced apical constriction provoked the folding of epithelial cell colonies on soft gels. Its application to murine and human neural organoids led to thickening of neuroepithelia, apical lumen reduction in optic vesicles, and flattening in neuroectodermal tissues. These results show that spatiotemporal control of apical constriction can trigger several types of 3D deformation depending on the initial tissue context.

## INTRODUCTION

Morphogenesis is the process by which cells organize to form 3D tissues and organs. The study of developing embryos has identified cell-level mechanisms that need to be coordinated to achieve morphogenesis. However, it is difficult to test the sufficiency of a mechanism to cause a specific change in tissue structure and to study feedback between multiple mechanisms during complex embryogenesis. As a solution, the field of synthetic morphology^1^ or synthetic developmental biology^2–6^ proposes to reconstitute morphogenetic events in vitro by gaining control of the constituent cell-level mechanisms.

Apical constriction, a process by which a cell actively reduces its apical surface, is necessary for the formation of numerous curved structures in metazoan embryos^7,8^. The driving force of apical constriction is actomyosin contraction, which is often triggered by activation of the Rho-ROCK pathway on the apical side. Because apical constriction occurs in specific stages and areas of the developing embryo, reconstituting curved tissues requires tools to control cellular contractility in space and time. Optogenetics is a powerful methodology to gain spatiotemporal control of biological processes from the molecular to the multicellular level^9–17^. Izquierdo and colleagues employed optogenetics to recruit RhoGEF to the plasma membrane in *Drosophila* embryos. Selective optogenetic activation on the apical side of dorsal cells led to tissue invagination, demonstrating that apical constriction is sufficient to induce deformation in that context^18^. Similar tools have been developed to spatiotemporally increase or reduce contractility in mammalian cells, mainly through recruiting RhoGEF, RhoA or myosin regulators to the plasma membrane. The approach has been effectively used to study mechanotransduction^19^, cell junction remodeling^20^, cytoskeletal dynamics^21^ and cytokinesis^22^. However, in mammalian tissues this approach has been mainly applied to study cell-level events, and its application to induce morphogenesis in complex tissue shapes would require very precise patterns of illumination. Therefore, there is still a lack of tools to manipulate 3D tissue deformation and reconstitute mammalian morphogenesis. In addition, the recent development of organoids, stem cell-derived 3D structures^23–25^, offers unique opportunities to study the interplay between tissue shape and function in vitro. However, the manipulation of organoid shape with optogenetic tools remains unexplored.

In this study, we present a new optogenetic tool that achieves control of apical constriction in mammalian cells, inducing multiple types of 3D tissue deformation. The tool is based on Shroom3, a key regulator of apical constriction necessary for several morphogenetic processes in vertebrates, including neural tube closure, lens placode invagination, and morphogenesis of gut and kidney^26–30^. The optogenetic version of Shroom3, OptoShroom3, is capable of fast activation and deactivation of apical constriction at the cell level. We demonstrate that an increase in apical tension causes tissue folding, thickening, flattening, and lumen shrinkage in epithelial cell sheets and neural organoids.

## RESULTS

To achieve spatiotemporal control over apical constriction in mammalian tissues, we created an optogenetic version of Shroom3. Shroom3 causes apical constriction by recruiting ROCK to apical junctions^31^. The Shroom Domain 1 (SD1) of Shroom3 is an actin-binding motif responsible for the apical localization^26,32,33^, whereas the SD2 is necessary for the binding to ROCK^31^ (figure 1a). The SD1 and SD2 domains are shown to function independently^26,34^. Therefore, we hypothesized that these domains could be split into two constructs and that the protein functionality could be restored through light-induced binding of the iLID-SspB optogenetic pair^35^. After testing multiple domain combinations, we found that the N-terminal Shroom3 fused with iLID (hereafter called NShroom3-iLID) and the C-terminal Shroom3 fused with SspB (SspB-CShroom3) function as an optogenetic split-version of Shroom3 (OptoShroom3) (figure 1b). GFP-NShroom3-iLID localized similarly to Shroom3 (figure 1c left; supp. figure 1) to the apical junctions of Madin-Darby Canine Kidney (MDCK) cells. By contrast, SspB-mCherry-CShroom3 acquired apical localization only upon seconds of blue light illumination (figure 1c right).

**Figure 1.**
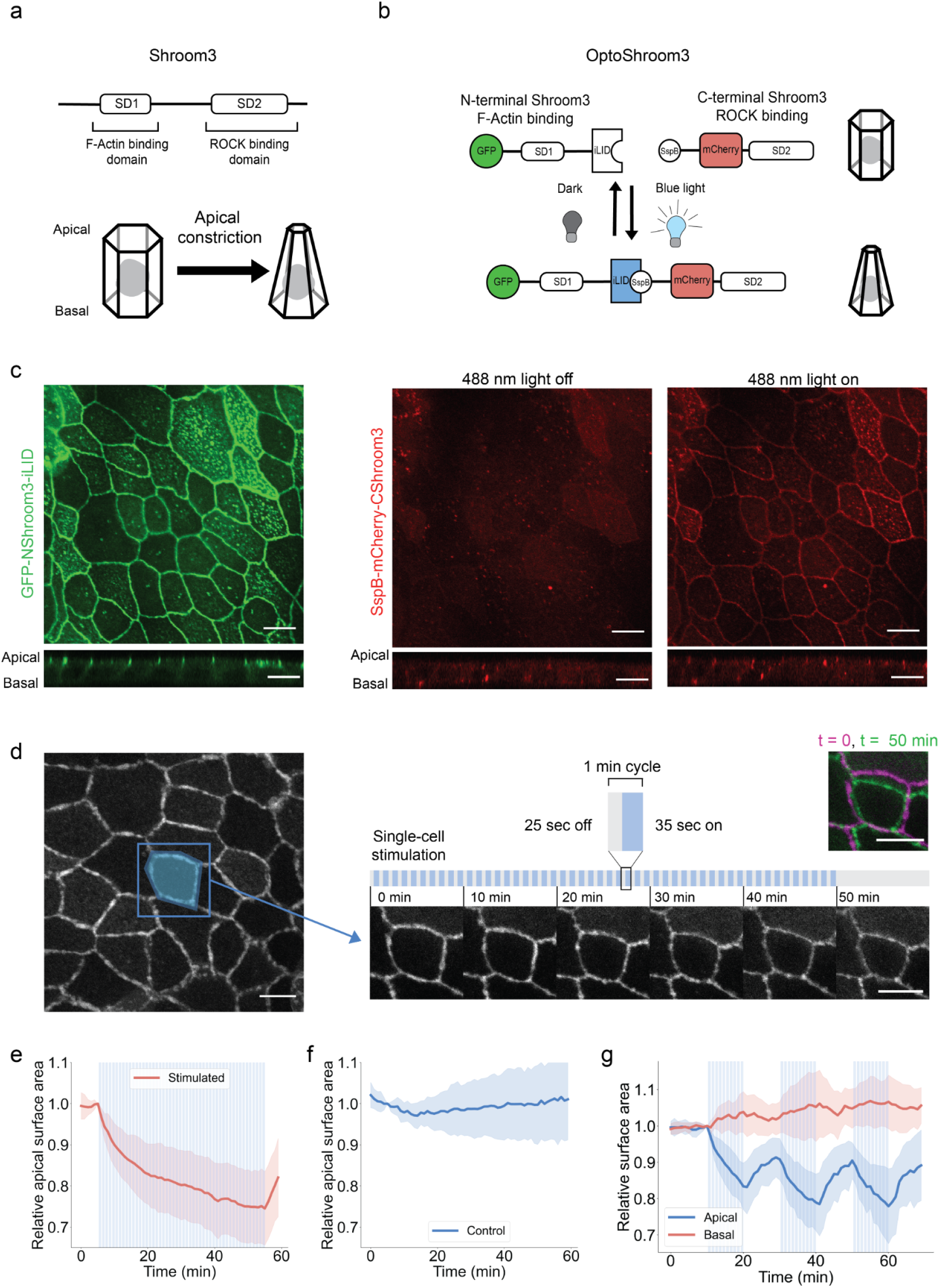
Characterization of OptoShroom3-induced apical constriction. **a**, Protein structure of wild-type Shroom3 and scheme of apical constriction. **b**, Design of OptoShroom3 constructs that dimerize upon blue light stimulation. **c**, MDCK cells expressing GFP-NShroom3-iLID and SspB-mCherry-CShroom3. Top: x-y apical slice, bottom: x-z lateral slice. Scale bar = 10 μm. **d**, Single-cell stimulation cycles and representative images from constriction experiments, apical slice. Stimulation area was designed as the initial apical area of the cell. iRFP-CAAX signal. Top right pane shows a color-coded compounded image comparing the start and end of stimulation. Scale bar = 10 μm. **e and f**, Quantification of the apical area in stimulated (N = 17) and non-stimulated (N = 16) cells. The area was normalized to the last measurement before stimulation (t = 5 min). Avg ± sd. **g**, Quantification of apical and basal areas during 3 periods of 10 min of stimulation and rest (N = 8, avg ± sd). The basal slice was defined as approximately 5 μm below the apical slice. The areas were normalized to the last measurement before stimulation (t = 10 min).

To test the ability of OptoShroom3 to induce apical constriction, we stimulated a single cell in an MDCK monolayer stably expressing OptoShroom3 constructs (figure 1d). The 1-minute illumination cycle was designed by taking into account the reported half-life of iLID-SspB binding, which is less than 1 minute^35^. The apical surface area showed a rapid decrease during the first minutes of stimulation, and constriction gradually decelerated, achieving a 25.4 ± 8.9% reduction of the original area within 50 minutes (figure 1d-f; supp. video 1). The apical surface started to increase only 1 minute after the end of stimulation, implying the fast unbinding of the two components and the deactivation of actomyosin contraction (figure 1e). These constriction dynamics are similar in shape and duration to those observed by Cavanaugh and colleagues using RhoGEF recruitment for cell-junction shortening in mammalian cells^20^. We then tested whether OptoShroom3 could be repeatedly activated and deactivated by concatenating several stimulation-rest periods (figure 1g; supp. video 2). The apical surface measurement indeed displayed periodic constrictions and fast activation-deactivation responses (less than 1 min each). By contrast, the basal surface did not show constriction upon stimulation but displayed a slightly increasing tendency during the stimulation phases. We reasoned that the subtle increase on the basal side could be due to membrane and cytoplasmic influx from the apical side, similarly to what has been described during *Drosophila* ventral furrow invagination^36^. These measurements collectively demonstrate that OptoShroom3 effectively reduces the apical-basal ratio of a stimulated cell (supp. figure 2) and has fast activation and deactivation kinetics.

Having measured the dynamics and functionality of OptoShroom3 in individually stimulated cells, we stimulated a larger region in a cell sheet to study the tissue-level effects. A confluent MDCK monolayer was formed on a thick collagen gel to provide an environment more permissive for deformations. We found that sustained photostimulation of a group of cells inside an illumination square (average 12 cells, figure 2a) caused cellular displacements towards the stimulated area on the apical side, whereas displacements in the opposite direction were observed after the end of stimulation (supp. video 3). Particle image velocimetry (PIV) analyses on the apical slices displayed a peak in inward displacements 2 minutes after stimulation, which gradually disappeared within the next 1 hour (figure 2b, c, d). A second peak of similar magnitude was observed in the outward direction immediately after the stimulation ended (figure 2d). Vectors showing the highest velocity, both upon start and end of stimulation, were those adjacent to the stimulated area (figure 2d, adjacent area). The summation of measured vectors along time accounted for a displacement of 1.31 ± 0.63 μm in the adjacent area (figure 2e). This result indicates that OptoShroom3 causes the largest deformation at the border between constricting cells and non-stimulated cells. Similar but smaller displacements were measured for the stimulated and outer area (0.84 ± 0.37 and 0.67 ± 0.29 μm, respectively). The highest displacement was achieved at 0-5 μm from the stimulated area, and displacement was observed up to 60 μm away from the stimulated area (supp. fig 3). Next, we produced a more complex deformation pattern through the simultaneous stimulation of two areas (figure 2f; supp. video 5). PIV analyses visualized tissue displacements with the shape of the stimulated areas (figure 2g). These results demonstrate that OptoShroom3 can induce 2D tissue-level deformations by pulling the apical surface of cells that are adjacent to stimulated areas.

**Figure 2.**
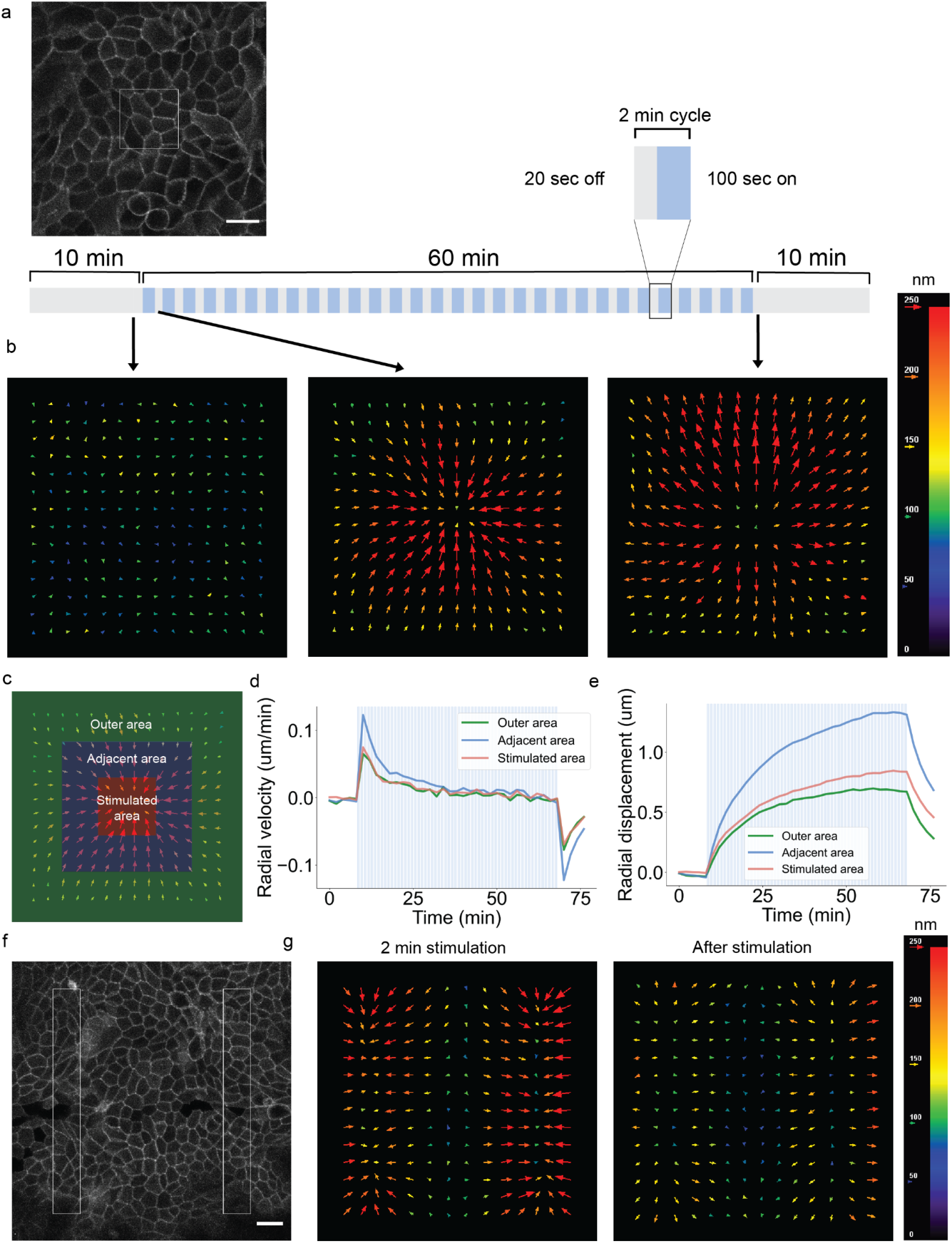
OptoShroom3-induced apical displacements on confluent monolayers. **a**, Apical slice of iRFP-CAAX signal of MDCK cells on a thick collagen gel. The white box marks the stimulated area. Scale bar = 20 μm. **b**, Averaged PIV analysis on apical slices before, during (2 min), and after stimulation in a 35.3 × 35.3 μm area (N = 6). **c**, Division of PIV frame in three areas for analysis, stimulated, adjacent, and outer areas. **d**, Temporal evolution of radial vector components averaged by area. Positive vectors represent movements towards the center of stimulation (N = 6, avg). **e**, Integration of vectors averaged by area (N = 6, avg). **f**, Apical slice of iRFP-CAAX signal of MDCK cells on a collagen gel. The white rectangles mark the stimulated areas. Scale bar = 20 μm. **g**, Averaged PIV analysis on apical slice during (2 min) and after stimulation of two 20 × 170 μm areas (N = 7).

To further assess if OptoShroom3 can induce 3D tissue morphogenesis in MDCK cells, we used matrigel, which is a well-known soft deformable gel. However, MDCK cells do not form a flat monolayer on a thick matrigel bed. We found that treatment of matrigel with acetic acid enabled the formation of flat MDCK monolayer colonies (supp. figure 4; figure 3a). Photostimulation of a whole colony in this setting caused folding of the MDCK cell sheet in 24 hours (figure 3b; supp. video 5). This time course resembles those of mammalian in vivo developmental folding processes, such as neural tube- and gut-folding, which require 1 or more days depending on the species^37–39^. Measurements of colony skeletons from x-z sections displayed an increase in average curvatures of stimulated colonies, compared with the unchanged curvatures of non-stimulated colonies (figure 3c, d). The curvature did not homogeneously increase in folding colonies, and the peripheral cells were displaced first towards the center of the colony, deforming and detaching from the substrate (figure 3b; supp. figure 5). Consistent with the increase in curvature, the projected area of stimulated colonies showed a decrease upon stimulation, reflecting the induced contractility (figure 3e, f). These measurements suggest that apical constriction changes the cell shape to pyramidal, inducing upward folding of a monolayer colony on a soft gel. To further test the controllability of the cell-sheet folding, we chose colonies with elongated shapes and stimulated a restricted area of a cell colony. Although the success of this approach depended on the initial shape of the colony, the spatial illumination led to the coiling and retraction of the stimulated area (figure 3g; supp. video 6). These results show that OptoShroom3 can provoke tissue curvatures and folds in a spatiotemporal manner.

**Figure 3.**
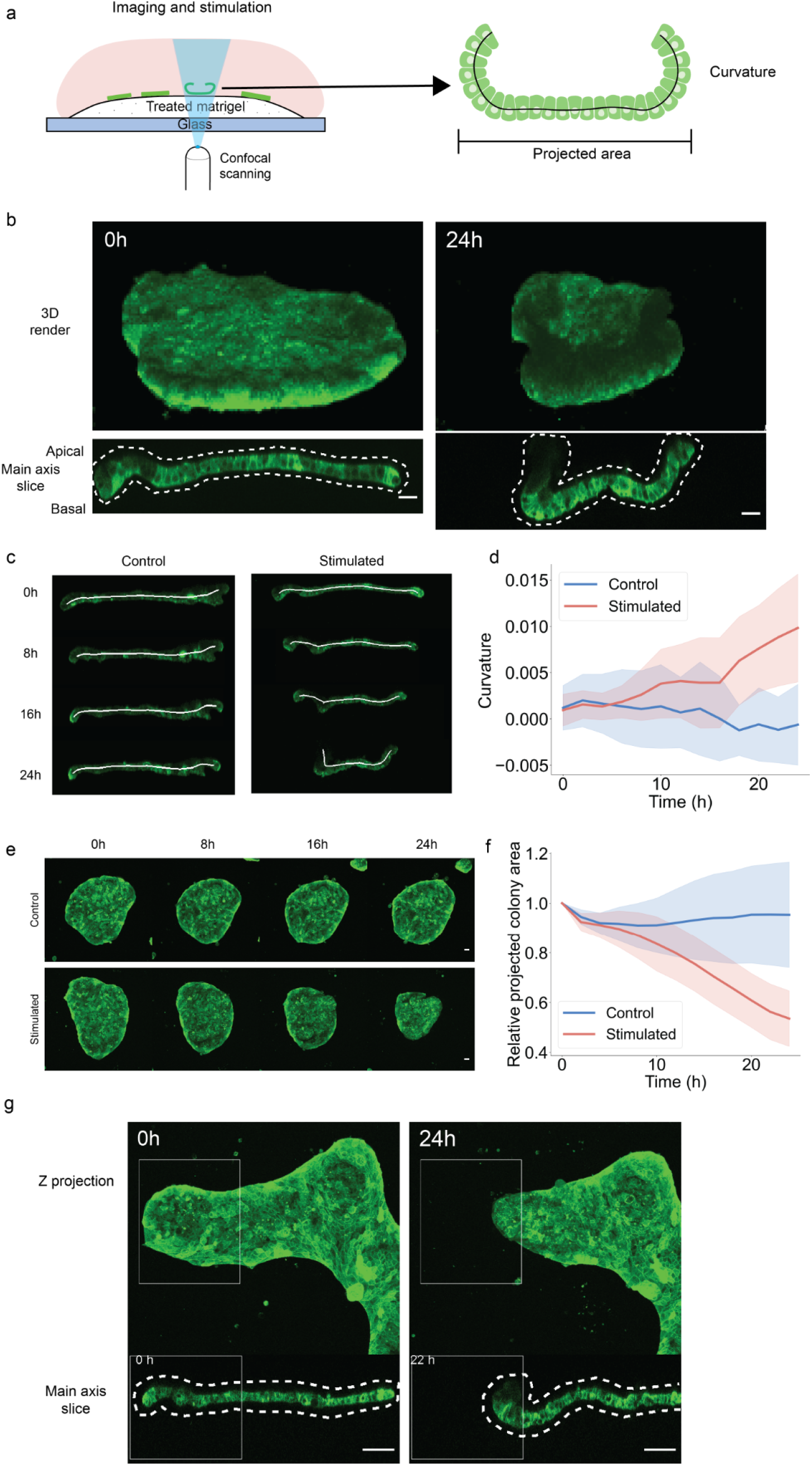
OptoShroom3-induced colony folding. **a**, Protocol for the formation of flat MDCK monolayers in a thick matrigel bed using acetic acid incubation (left) and scheme of measured parameters (right). **b**, 3D render and higher resolution lateral slices of representative colony folding before and after 24 h of stimulation. GFP-NShroom3-iLID expression. Scale bar = 20 μm. **c**, Curvature measurements of representative examples of non-stimulated (left) and stimulated (right) colonies. GFP-NShroom3-iLID expression. **d**, Quantification of changes in average curvature of stimulated and non-stimulated colonies. Concave (apical) curvature was set as positive (Nstim = 7, Ncont = 7, avg ± sd). **e**, Projected areas of representative non-stimulated (top) and stimulated (bottom) MDCK colonies for 24 h. GFP-NShroom3-iLID expression. Scale bar = 20 μm. **f**, Quantification of relative changes in the projected area of stimulated and non-stimulated colonies (Nstim = 7, Ncont = 7, avg ± sd). **g**, Z-projection and re-slices of the selectively stimulated colony, the white rectangle marks stimulated regions. GFP-NShroom3-iLID expression. Scale bar = 50 μm.

To alter mammalian tissue structure in more complex systems, we further tested the application of OptoShroom3 in mouse optic vesicle organoids. Here we denominate optic vesicle organoids to the earlier stages of optic cup organoids^40^, which are known to develop polarized neuroepithelia that forms optic vesicles. Optic cups have been demonstrated to form through the regulation of apical constriction^40,41^. Towards this aim, we prepared a mouse embryonic stem (mES) cell line stably expressing OptoShroom3 and created optic vesicle organoids (figure 4a). GFP-NShroom3-iLID changed its localization along with organoid differentiation, from a diffused signal (days 1-3) to a concentrated signal on the apical side of the neuroepithelium (day 4-) (supp. figure 6). It is important to mention that the inner surface of optic vesicle organoids is apical. We considered the change in localization an indication of the proper differentiation of the neuroepithelium. Photostimulation of OptoShroom3 in the earlier stages of a neuroepithelium (days 4-8) caused translocation of SspB-mCherry-CShroom3 and a 14.3 ± 7.4% increase in the thickness of the epithelial layer (supp. video 7; figure 4b, c). The stimulation of a whole optic vesicle (days 6-8) caused a decrease in the size of the apical lumen (figure 4d, e; supp. video 8). The longest diameter of the lumen displayed a 9 ± 5.8% decrease after ∼1 hour of continuous stimulation (figure 4f, g). We reasoned that apical constriction made individual cells taller, effectively reducing the apical lumen size.

**Figure 4.**
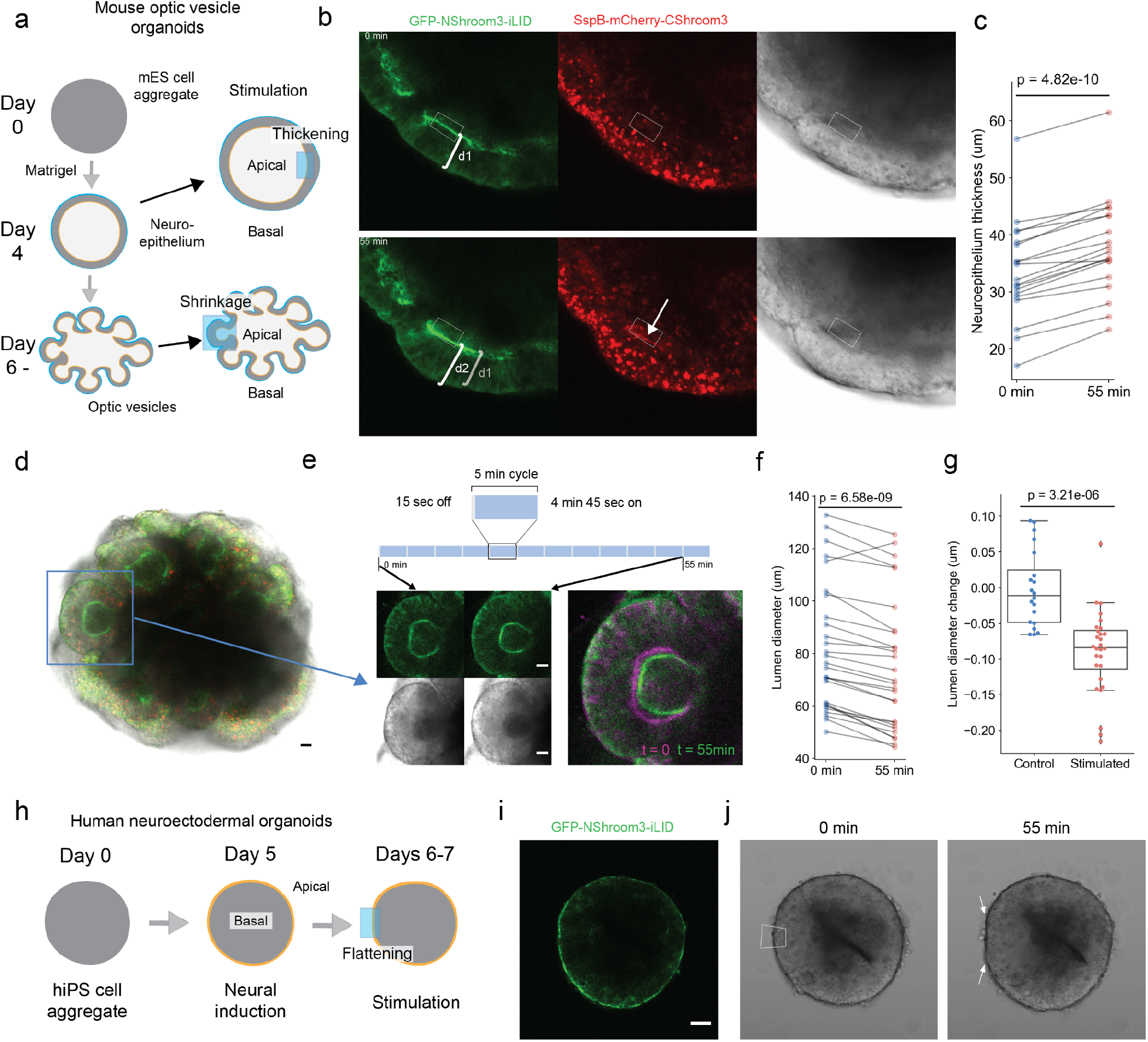
OptoShroom3-induced tissue deformation in mouse and human neural organoids. **a**, The formation and stages of mouse optic vesicle organoids. **b**, Representative example of stimulation of neuroepithelium. Arrow indicates the translocation of SspB-mCherry-CShroom3. **c**, Thickness measurements of neuroepithelia before and after stimulation (days 4-8) (N = 19, paired t-test). **d**, Representative example of optic vesicle organoid on day 8. **e**, Stimulation cycles and representative example of 488 nm stimulation of an optic vesicle showing transmitted light and GFP-NShroom3-iLID signal. Right panel shows a color-coded compounded image comparing the start and end of stimulation. **f**, Measurements of lumen diameter of optic vesicles before and after OptoShroom3 stimulation (days 6-8) (N = 27, paired t-test). **g**, Quantification of the lumen diameter change for control (non-stimulated) and stimulated samples after 55 min (Ncont = 17, Nstim = 27, student’s t-test). **h**, Human neuroectodermal organoid formation stages. **i**, GFP-NShroom3-iLID expression on a neuroectodermal organoid, day 6. **j**, Representative example of OptoShroom3 induced flattening through local stimulation of neuroectodermal organoids. Scale bar = 50 μm.

To test if the reversed apicobasal polarity leads to a different type of deformation, we utilized human neuroectodermal organoids. Neuroectodermal organoids were formed from human induced pluripotent stem (hiPS) cells stably expressing OptoShroom3, following the first steps of cerebral organoid formation^42^ (figure 4h). Contrary to optic vesicle organoids, neuroectodermal organoids show outer apical polarity before matrigel addition, and therefore exhibit a convex apical side. Accordingly, we observed that GFP-NShroom3-iLID localized on the outer surface of the organoid upon neural induction (days 6-7) (figure 4i). Selective stimulation of a border of neuroectodermal organoids provoked flattening of the area in less than 1 hour (figure 4j), reducing the initial curvature and provoking inward tissue displacement (supp. video 9). These results demonstrate that OptoShroom3 can manipulate morphologies of organoids, including epithelial thickening, lumen reduction, and flattening. The outcome of deformation depends on initial tissue geometry, apicobasal polarity, and the forces to which the tissue is already subjected to.

## DISCUSSION

In this study, we have developed a novel optogenetic tool to control apical constriction in mammalian tissues. The tool induced a quick reduction of the apical cell surface, and the stimulation of groups of cells provoked collective displacements in the apical side of adjacent cells. Apical constriction in epithelial colonies on soft gels induced the folding of cell sheets. In organoids, apical constriction thickened the neuroepithelium and reduced the inner lumen when the inner surface was apical. By contrast, it flattened the tissue when the outer surface of organoids was apical.

These results illustrate how the induction of apical constriction in different contexts can lead to different changes in tissue structure. While the cuboidal epithelium formed by MDCK cells acquired curvature and underwent folding, the neuroepithelium of optic vesicle organoids thickened in less than 1 hour without necessarily acquiring curvature. A possible explanation for these differences is the specific architecture and availability of cytoskeletal components of each cell type, which could potentially condition the cell shape changes that can be achieved. In addition, the function of Shroom3 may be more complex than the induction of actomyosin constriction. Although apical constriction could itself be enough to explain cell elongation through cell volume conservation^43,44^, Shroom3 has also been reported to induce a redistribution of γ-tubulin, a microtubule regulator, to cause cell elongation along the apicobasal axis^45^.

OptoShroom3 joins the expanding set of optogenetic tools capable of inducing actomyosin constriction. While pre-existing tools have mainly used plasma membrane recruitment of RhoGEF or RhoA factors for the activation of actomyosin constriction^18–21^, the recruitment of OptoShroom3 is specific to the apical junctions of epithelial cells. Therefore, OptoShroom3 does not require subcellular precision of stimulation with multi-photon microscopy to induce apical constriction. If a whole cell is stimulated, SspB-mCherry-CShroom3 will be recruited to the apical side and induce constriction in that area only. These features highlight OptoShroom3 as a new tool to externally manipulate and control tissue shapes and to study tissue mechanics^46–48^ and mechanotransduction^49,50^.

The rapid activation-deactivation cycle of OptoShroom3 can be an advantage when studying dynamic and fast processes. Although this may be considered a limitation when planning long-term experiments, continuous stimulation did not cause visible phototoxicity in the 24-hour folding experiments. A potential way to reduce stimulation periods would be to take advantage of mutant versions of LOV2 domain of iLID known to increase the half-life of its binding to SspB^51^.

In the long term, novel tools that can induce morphological changes will be necessary not only to study in vivo morphogenesis but also to reproduce the morphogenetic processes in vitro. We expect that OptoShroom3 will provide a new avenue towards understanding how tissue shape and mechanotransduction affect cell fate and differentiation. Organoids, given their inherent complexity and accessibility, stand out as perfect candidates for the application of these new “morphogenetic tools” to further study the interplay between tissue shape and function.

## MATERIALS AND METHODS

### Cloning and construction of optogenetic tool

For OptoShroom3 design, mouse *Shroom3* gene was used, which was a gift from T. Nishimura, from M. Takeichi lab. *Shroom3b* isoform (originally named *ShrmS*^26^), which presents an N-terminal deletion of 177 aa that removes a PDZ domain, was used for the construction. Construction of OptoShroom3 was carried out using iLID and Sspb genes from the optogenetic vector library published by Tichy and colleagues^52^, which was a gift from H. Janovjack lab. OptoShroom3 sequences can be found in supplementary table 1. OptoShroom3 constructs GFP-NShroom3-iLID and SspB-mCherry-CShroom3 have been submitted to Addgene (catalogue numbers: 170976 and 170977 respectively). iRFP-CAAX and GFP-CAAX sequences were prepared through traditional cloning methods. iRFP gene was acquired from addgene (piRFP, plasmid #31857). For stable construct expression, CAG promoter was used for OptoShroom3 and GFP-CAAX and Ef1α promoter for iRFP-CAAX.

### Cell culture

MDCK cell line (MDCKII) was a gift from M. Murata lab. MDCK cells were maintained in Dulbecco’s Minimal Essential Medium (DMEM) with high glucose, GlutaMax, and pyruvate (Gibco, 31966-021), supplemented with 10% fetal bovine serum, penicillin, and streptomycin (100 U/ml and 100 μg/ml, Gibco). Cells were split every 2-3 days through 9 minutes incubation with EDTA 50mM pH=7.4 and 3 minutes of 0.25% trypsin (Gibco).

Mouse ES cell line (EB5, Rx-GFP) was described previously^40^. mES cells were maintained on gelatin-coated dishes with DMEM supplemented with 15% fetal bovine serum, nonessential amino acids, GlutaMax, sodium pyruvate (Gibco, 11360-039), β-mercaptoethanol (0.1 mM), LIF (1500 U/ml), CHIR99021 (3 µM), and PD0325901 (1 µM). The 2i medium was used to increase the number of optic vesicles per organoid. Cells were split every 2-3 days with 0.25% trypsin.

Human iPS cell line ((IMR90)-4) was purchased from WiCell. hiPS cells were maintained in StemFlex media (Gibco) on matrigel-coated dishes. DMEM-F12 (Gibco) with 0.013% matrigel (Corning #356231) was used for coating for 30 minutes at 37°C. Cells were split every 4-5 days with Accutase (STEMCELL) and StemFlex media was supplemented with ROCK inhibitor Y-27632 (10 µM) during the first day after splitting.

For MDCK and mES cells, OptoShroom3, iRFP-CAAX, and GFP-CAAX constructs were stably transfected using PiggyBac system^53^ and lipofectamine. For hiPS cells, PiggyBac plasmids were introduced using Amaxa Nucleofector (Lonza). After antibiotic selection, stable clones were picked for every cell line. All cell lines were cultured at 37ºC in a humidified incubator with 5% CO2 and regularly tested for mycoplasma contamination.

### Imaging and stimulation of optogenetic tool

All images were acquired using Olympus FV3000 confocal microscope at 37ºC and 5% CO2. Stimulation of OptoShroom3 in all samples was performed using stimulation mode and 0.01-0.03% power of 488 nm laser. UPLSAPO40XS and UPLSAPO30XS objectives were used for MDCK imaging and stimulation, and UPLSAPO10X2 was used for organoid imaging and stimulation.

### Image analysis

Apical and basal area measurements were conducted on a custom-made python pipeline, applying watershed segmentation to iRFP-CAAX confocal stacks of MDCK cells on glass-bottom dishes. Basal stacks were selected as approximately 5 µm below apical. Projected area and skeletons for curvature measurements of folding MDCK colonies were obtained from confocal stacks using a custom-made python segmentation pipeline.

### Curvature measurement

Local curvature was defined as the inverse of the radius of the osculating circle to three points of a curve. Curvature was measured by calculating the osculating circle of three equally distanced points (20 pixels distance) in the skeleton, equivalent to calculating the inverse of the radius of a circumscribed circle around a triangle:

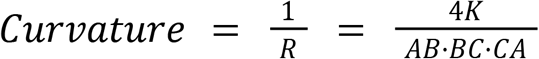

Being R the radius of the osculating circle of points A, B, and C. AB, BC, and CA are the distance between the points. K is the area of the triangle formed by A, B, and C. Local curvatures were measured along the skeleton of the main axis and averaged for every time point to obtain a general curvature measurement. Curvature measurements in figure 3 are in units 1/pixel.

### PIV analysis

Particle Image Velocimetry analyses were conducted on iRFP-CAAX projections of apical slices of MDCK cells seeded in 100% collagen gels. For preparation, 50ul of 100% collagen I were added to a glass bottom dish and incubated at 37ºC for 30mins for polymerization. After incubation, 5 × 10^5^ OptoShroom3+iRFP-CAAX MDCK cells were seeded in 80 μl of DMEM medium. More media (2 ml) was added 1.5 hours after seeding. Confluent monolayers were stimulated 2 days after seeding using 0.03% power of 488 nm laser. Images were acquired every 2 minutes. PIV analysis was performed using ImageJ PIV analysis plugin developed by Q. Tseng. The cross-correlation method was used with 1 pass with an interrogation window size of 64 pixels. Batch analysis, classification of vectors by area, and experiment averaging were performed with a custom python pipeline.

### Tissue folding

For tissue folding assays, 20 μl of 100% matrigel (Corning #356234) were set on glass-bottom dishes and solidified at 37ºC for 10 minutes. Then, gels were incubated with 20 mM acetic acid (90 μl, 4.5 matrigel-acetic acid volume ratio) at 37ºC for 40 minutes. After incubation, gels were washed twice with PBS and 4 × 10^5^ MDCK cells were seeded in 80 μl of DMEM medium. More media (2 ml) was added 1.5 hours after seeding. These parameters were empirically obtained to make isolated flat MDCK colonies, which were stimulated 2 days after seeding.

Stimulated colonies were illuminated for 57 minutes every hour. The remaining 3 minutes were used for the acquisition of GFP-NShroom3-iLID signal in control colonies (non-stimulated colonies). We considered the impact of this period of illumination of control colonies and of lack of illumination of stimulated colonies negligible. Colonies that showed migratory behaviour were not considered for the analysis.

### Optic vesicle organoids

Optic vesicle organoids were prepared following SFEBq method^40^. mES cells expressing OptoShroom3 were dissociated with 0.25% trypsin and quickly reaggregated in differentiation medium, G-MEM supplemented with 1.5% knockout serum replacement, nonessential amino acids, sodium pyruvate, and β-mercaptoethanol (0.1 mM). 3000 cells were cultured in 100 µl media per well of a 96 well low attachment U bottom plate (Thermo). Matrigel (1%, Corning #356231) was added the next day.

Although the ES cells express the retinal marker Rx-GFP during the optic vesicle formation, the Rx-GFP expression is much lower than that of GFP-NShroom3-iLID and thus negligible. Measurements for neuroepithelial thickening and lumen reduction were manually performed using FIJI. Each measurement was performed three times and averaged.

### Neuroectodermal organoids

Neuroectodermal organoids were prepared using hiPS cells expressing OptoShroom3 following the first steps of cerebral organoid protocol^54^ with STEMdiff(tm) Cerebral Organoid Kit (STEMCELL Technologies, #08570). The modifications from commercial protocol were: 2000 cells were used as starting number, Y-27632 (10 µM) was added from day 0 to day 3, and the use of Accutase for cell disaggregation.

## ACKNOWLEDGMENTS

This work was supported by internal grants from RIKEN and EMBL; Grant-in-Aid for Scientific research (KAKENHI) programs from Japanese Ministry of Education, Culture, Sports, Science, and Technology (MEXT) (16H06170 and 18H04769 to M.E.); Research foundation for Opto-Science and Technology (to M.E.).

We are thankful to M. Matsuda for his help in cell-line preparation and discussions about OptoShroom3; to Y. Sasai for his help in optic cup organoids; to A. Malandrino and M. Marin for their helpful conversations on gel preparation and tissue folding; to L. Batlle and the Tissue Engineering unit from the Center for Genomic Regulation for their instruction and comments in organoid culture; to M. Lancaster and S. Benito-Kwiecinski for their instruction in the formation of neuroectodermal organoids and helpful suggestions; to X. Trepat and M. Gomez for their help in gel measurements and comments; to N. Montserrat, M. Gallo and E. Garreta for their support in experiment design and comments on the manuscript; to V. Ruprecht, J. Davies, S. de Renzis, M. Bosch, T. Matsui, J. Lázaro, M. Costanzo, and K. Aoki for their comments on the manuscript.

## SUPPLEMENTARY FIGURES

**Supplementary figure 1.**
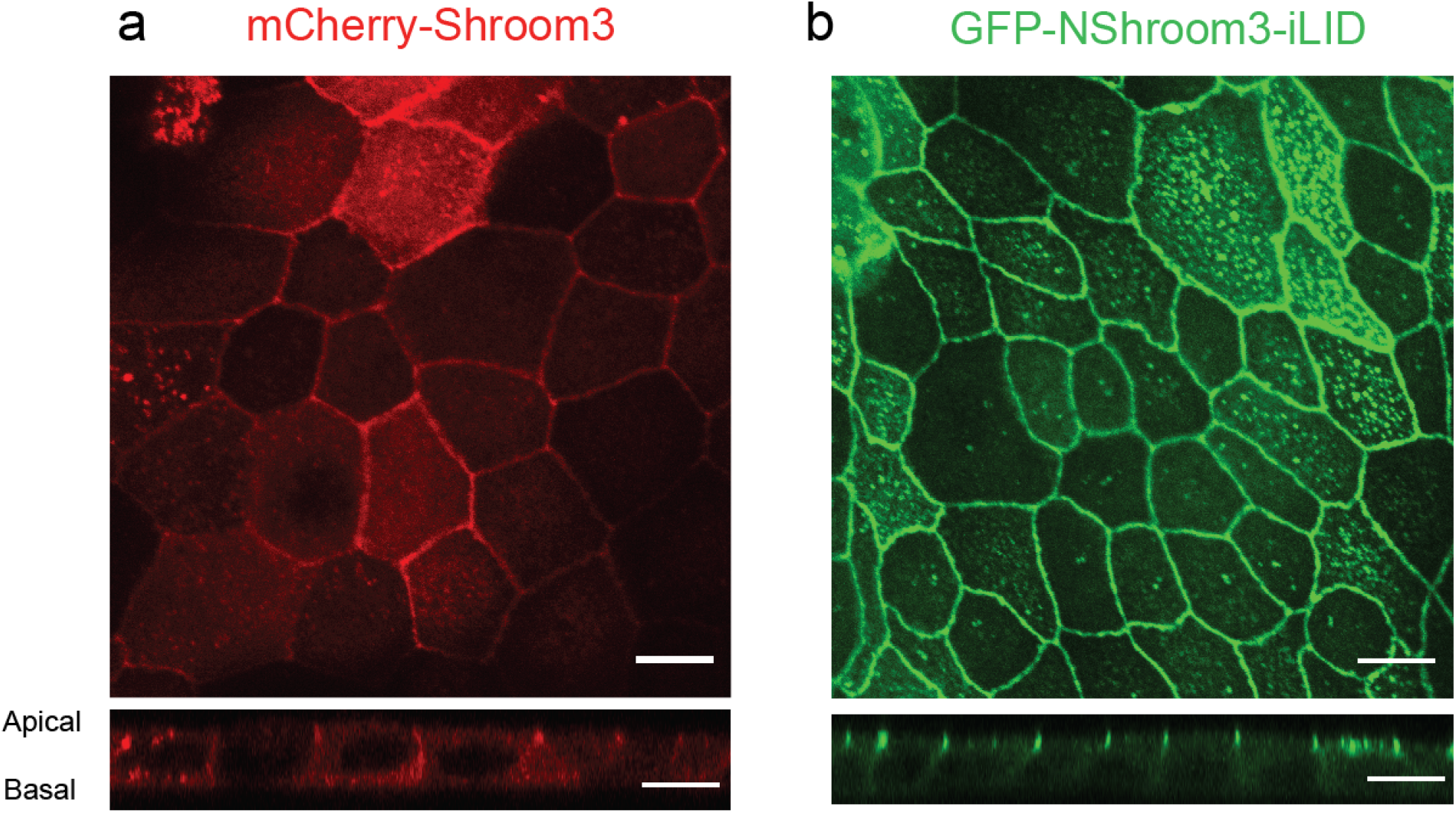
GFP-NShroom3-iLID localizes in the apical junctions, similarly to Shroom3. **a**, MDCK cells expressing mCherry-Shroom3. **b**, MDCK cells expressing GFP-NShroom3-iLID (same image as in figure 1c). Top: x-y apical slice, Bottom: x-z lateral slice. Scale bars = 10 μm.

**Supplementary figure 2.**
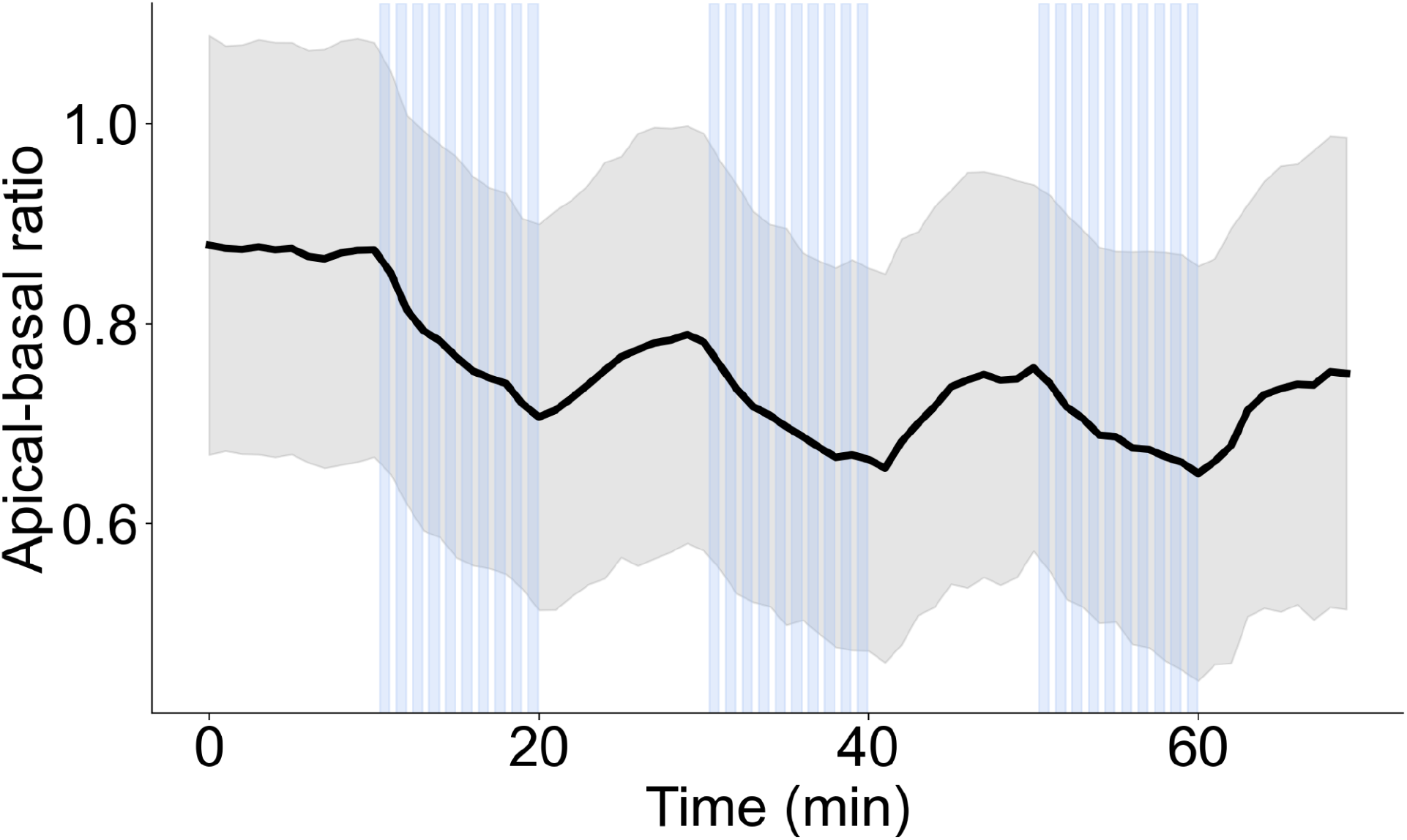
Apical-basal ratio reduction upon stimulation and partial recovery during non-stimulation phases. Calculated from data displayed in figure 1g, the ratio of apical and basal areas measured by segmentation of MDCK cells. Stimulation cycle: 25 sec image acquisition, 35 sec stimulation (blue). Repetition of 10 cycles of rest (no stimulation) and 10 cycles of stimulation (N = 8, avg ± sd).

**Supplementary figure 3.**
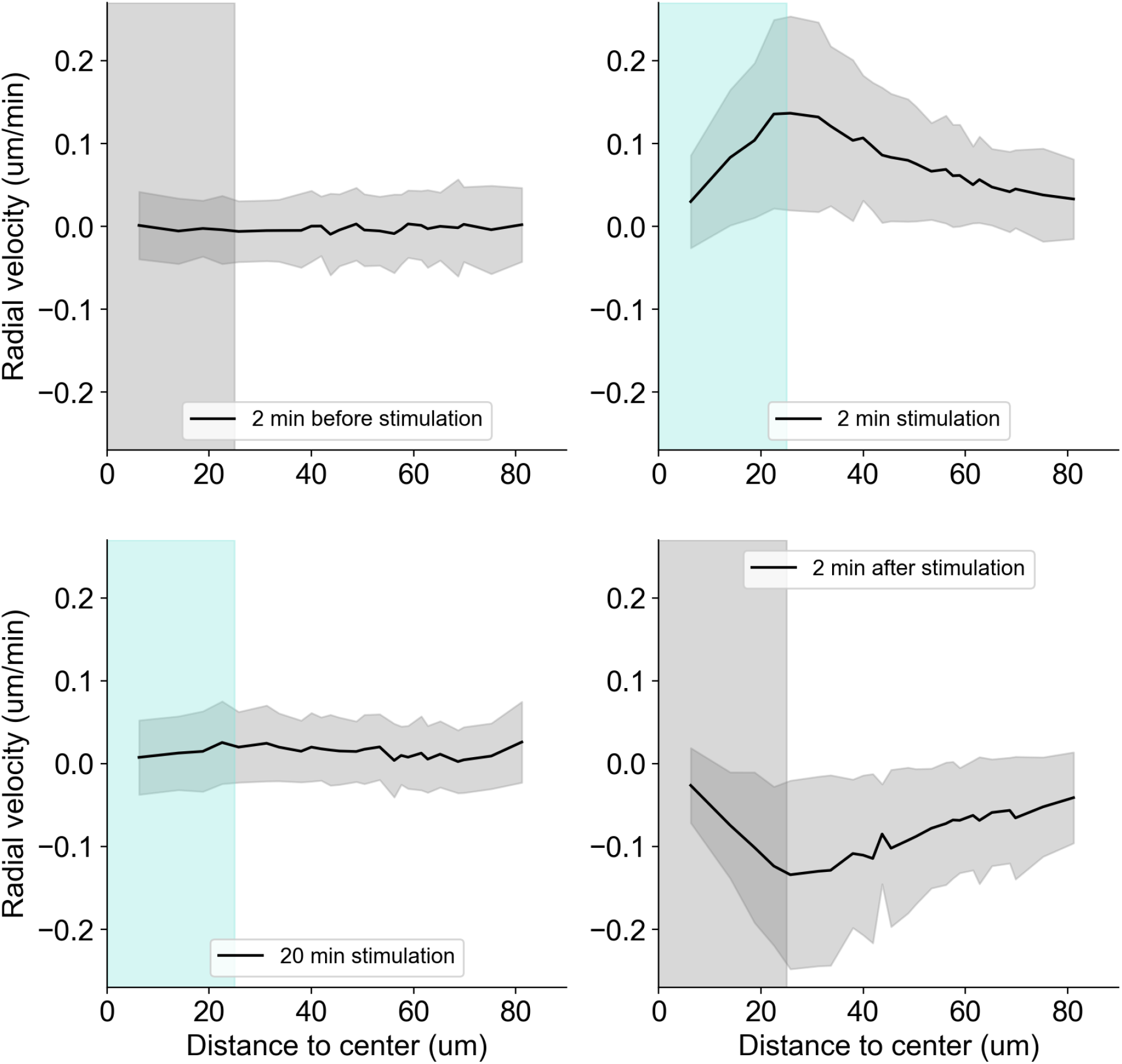
OptoShroom3 induced apical displacement averaged by distance to the center of stimulation. **a**,**b**,**c and d**, Average displacements caused by stimulation of a 35.3 × 35.3 μm square on confluent MDCK cells on 100% collagen gel before stimulation, after 2 minutes of stimulation, after 20 minutes of stimulation, and 2 minutes after the end of stimulation. Stimulated area displayed in blue as the distance from the center to the corner of the square (24.96 μm) (N = 6, avg ± sd) (Reanalysis of data from figure 2).

**Supplementary figure 4.**
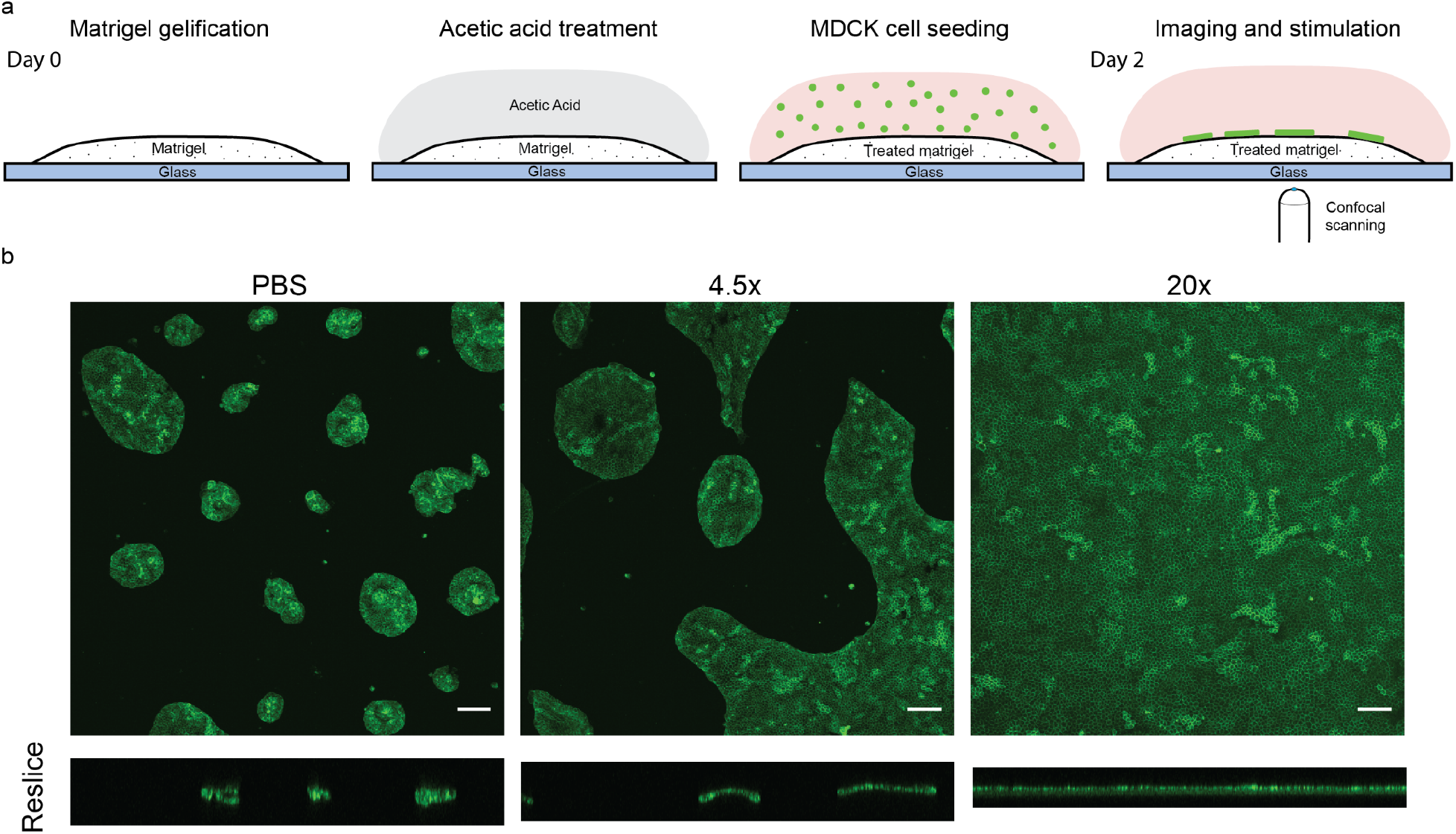
Acetic acid treatment on matrigel affects MDCK colony formation. **a**, Protocol to improve flat MDCK monolayer formation on matrigel settings. A thick matrigel gel was polymerized on a glass bottom dish, then incubated with 20 mM acetic acid before cell seeding. **b**, Comparison of MDCK colonies formed on untreated (PBS) matrigel with those formed on gels treated with two different volumes of acetic acid (acetic acid/matrigel = 4.5 and 20). Same number of cells was seeded on the three gels. GFP-CAAX signal.

**Supplementary figure 5.**
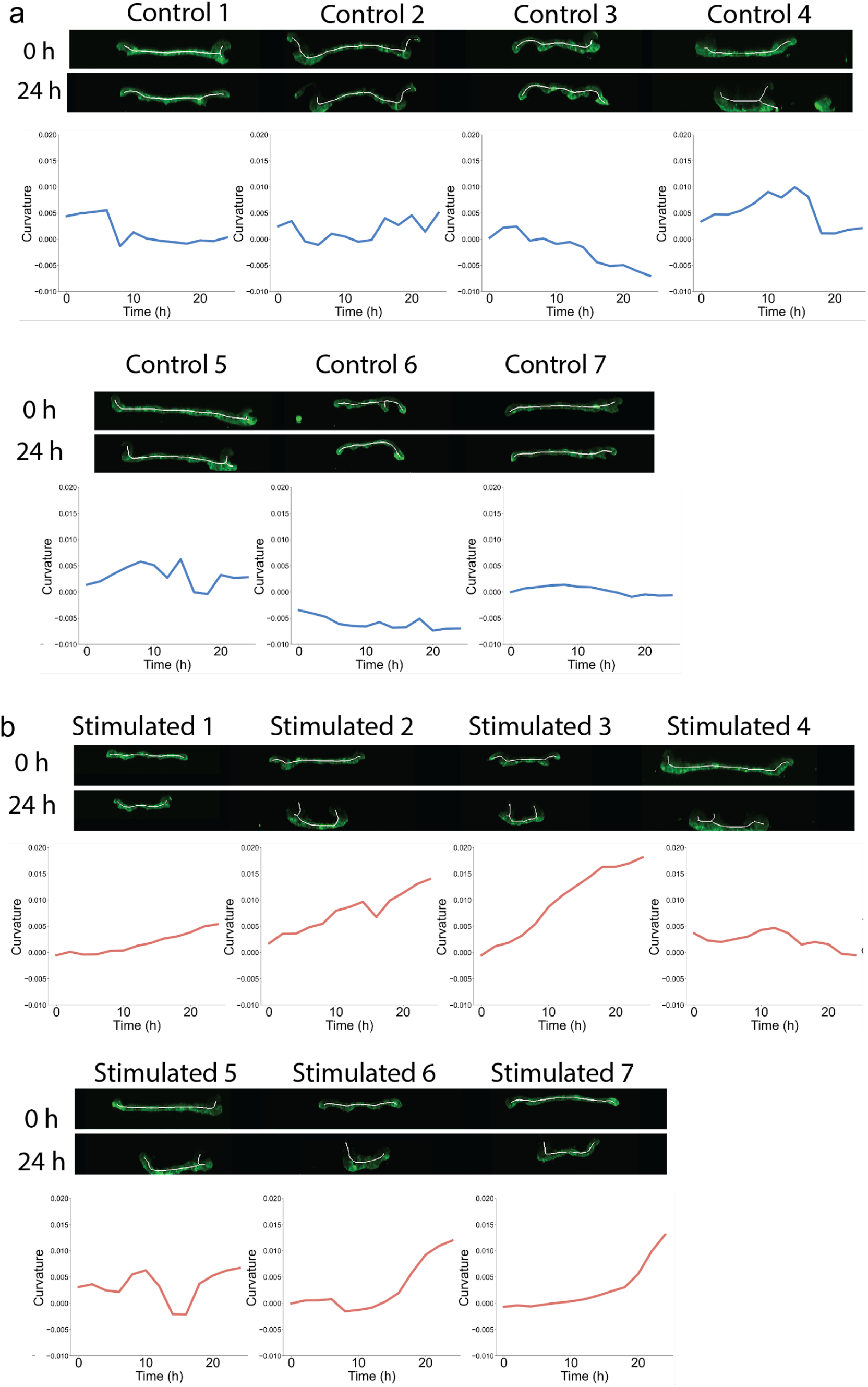
Summary of MDCK cell colony folding. **a, b**, Summary of average curvature measurements from centric line of control and stimulated colonies averaged in figure 3 measurements (Ncont = 7, Nstim = 7) (Raw data from figure 3. Control 7 and stimulated 7 are also displayed on figure 3).

**Supplementary figure 6.**
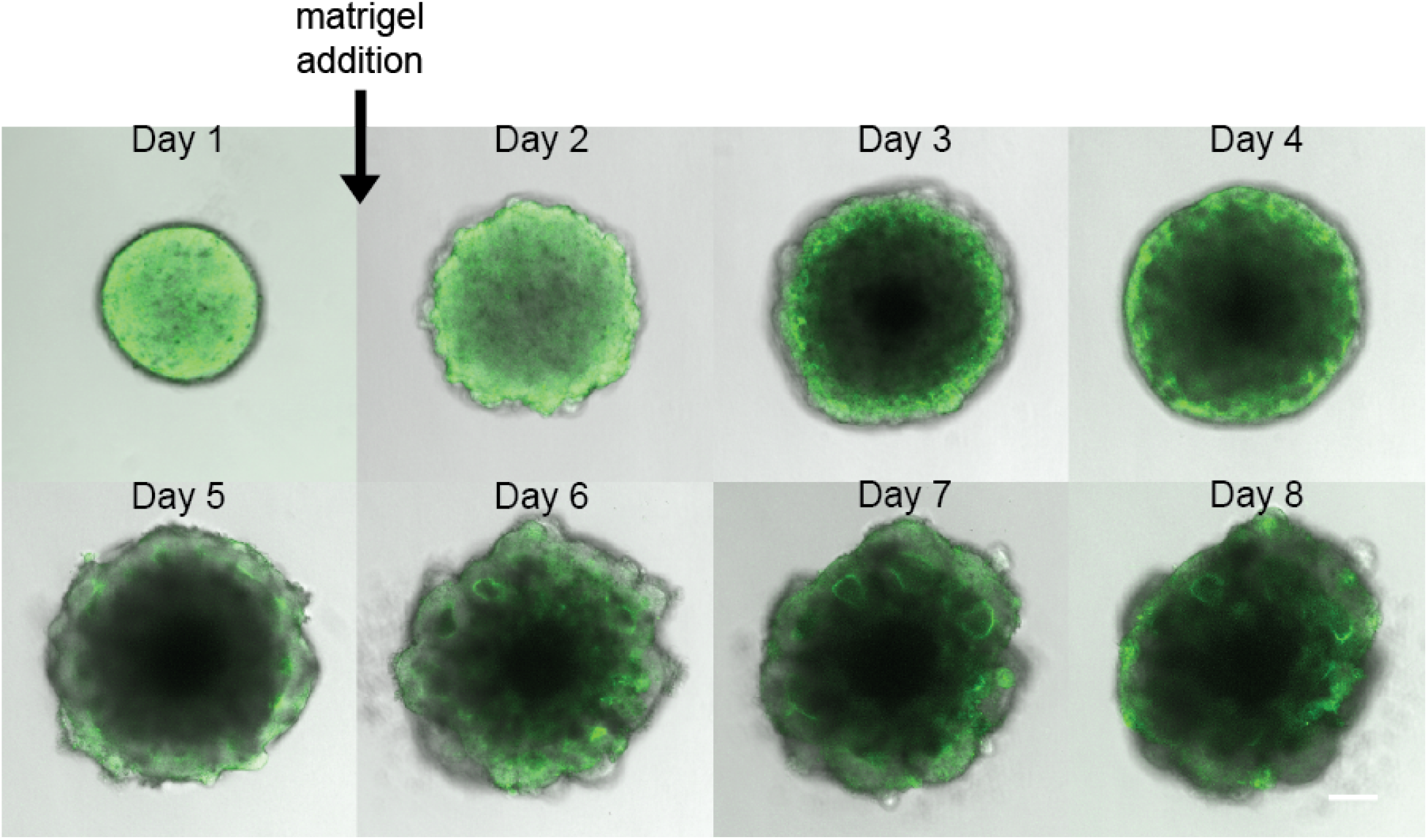
Optic vesicle organoid development and change in GFP-NShroom3-iLID localization. During the first 8 days of optic vesicle organoid formation, GFP-NShroom3-iLID localization changed from homogeneous distribution to a strong localization on the apical side of vesicles. Scale bar = 100 μm.

## SUPPLEMENTARY DATA

All supplementary videos are displayed 3 times.

**Supplementary video 1**. Translocation of SspB-mCherry-CShroom3 and apical constriction induced by stimulation of a single cell in an OptoShroom3-MDCK monolayer. At minute 20, the microscope was stopped for less than 1 minute to set more stimulation cycles, which caused a transient decrease in translocation.

**Supplementary video 2**. Repeated stimulation of a single cell in an OptoShroom3-MDCK monolayer. iRFP-CAAX. Apical and basal slices. 3 stimulation rounds.

**Supplementary video 3**. Stimulation of a group of cells in an OptoShroom3-MDCK monolayer. iRFP-CAAX.

**Supplementary video 4**. Simultaneous stimulation of two areas in an OptoShromm3-MDCK monolayer. iRFP-CAAX.

**Supplementary video 5**. Stimulation-induced folding of OptoShroom3-MDCK colonies. GFP-NShroom3-iLID.

**Supplementary video 6**. Retraction and coiling of cell sheets induced by selective stimulation of elongated OptoShroom3-MDCK colonies. Showing 5 different samples. GFP-NShroom3-iLID.

**Supplementary video 7**. Thickening of a neuroepithelium induced by selective stimulation of an OptoShroom3-optic vesicle organoid. GFP-NShroom3-iLID and Sspb-mCherry-CShroom3.

**Supplementary video 8**. Apical lumen reduction induced by selective stimulation of OptoShroom3-optic vesicle organoids. GFP-NShroom3-iLID signal.

**Supplementary video 9**. Flattening of outer membranes induced by selective stimulation of OptoShroom3-neuroectodermal organoids.

**Supplementary table 1**. Sequences of OptoShroom3 constructs.

## Notes

### Competing Interest Statement

The authors have declared no competing interest.

### Summary of Updates

Minor correction and updates regarding the genetic constructs (Fig 1 and method)

## REFERENCES

1. Davies, J. A. Synthetic morphology: prospects for engineered, self-constructing anatomies. J. Anat. 212, 707–719 (2008).

2. Ebrahimkhani, M. R. & Ebisuya, M. Synthetic developmental biology: build and control multicellular systems. Curr. Opin. Chem. Biol. 52, 9–15 (2019).

3. Toda, S., Brunger, J. M. & Lim, W. A. Synthetic development: learning to program multicellular self-organization. Curr. Opin. Syst. Biol. 14, 41–49 (2019).

4. Santorelli, M., Lam, C. & Morsut, L. Synthetic Development: building mammalian multicellular structures with artificial genetic programs. Curr. Opin. Biotechnol. 59, 130–140 (2019).

5. Elowitz, M. & Lim, W. A. Build life to understand it. Nature 468, 889–890 (2010).

6. Teague, B. P., Guye, P. & Weiss, R. Synthetic Morphogenesis. Cold Spring Harb. Perspect. Biol. 8, a023929 (2016).

7. Sawyer, J. M. et al. Apical Constriction: A Cell Shape Change that Can Drive Morphogenesis. Dev. Biol. 341, 5–19 (2010).

8. Martin, A. C. & Goldstein, B. Apical constriction: themes and variations on a cellular mechanism driving morphogenesis. Development 141, 1987–1998 (2014).

9. Baumschlager, A. & Khammash, M. Synthetic Biological Approaches for Optogenetics and Tools for Transcriptional Light-Control in Bacteria. Adv. Biol. n/a, 2000256.

10. Ruijgrok, P. V. et al. Optical control of fast and processive engineered myosins in vitro and in living cells. Nat. Chem. Biol. 1–9 (2021) doi:10.1038/s41589-021-00740-7.

11. Forlani, G. & Di Ventura, B. A light way for nuclear cell biologists. J. Biochem. (Tokyo) 169, 273–286 (2021).

12. Gagliardi, P. A. & Pertz, O. Developmental Erk Signaling Illuminated. Dev. Cell 48, 289–290 (2019).

13. Johnson, H. E. & Toettcher, J. E. Illuminating developmental biology with cellular optogenetics. Curr. Opin. Biotechnol. 52, 42–48 (2018).

14. Mumford, T. R., Roth, L. & Bugaj, L. J. Reverse and forward engineering multicellular structures with optogenetics. Curr. Opin. Biomed. Eng. 16, 61–71 (2020).

15. Hartmann, J., Krueger, D. & De Renzis, S. Using optogenetics to tackle systems-level questions of multicellular morphogenesis. Curr. Opin. Cell Biol. 66, 19–27 (2020).

16. Zhang, Z. et al. Optogenetic manipulation of cellular communication using engineered myosin motors. Nat. Cell Biol. 23, 198–208 (2021).

17. Tischer, D. & Weiner, O. D. Illuminating cell signalling with optogenetic tools. Nat. Rev. Mol. Cell Biol. 15, 551–558 (2014).

18. Izquierdo, E., Quinkler, T. & De Renzis, S. Guided morphogenesis through optogenetic activation of Rho signalling during early Drosophila embryogenesis. Nat. Commun. 9, 2366 (2018).

19. Valon, L., Marín-Llauradó, A., Wyatt, T., Charras, G. & Trepat, X. Optogenetic control of cellular forces and mechanotransduction. Nat. Commun. 8, 14396 (2017).

20. Cavanaugh, K. E., Staddon, M. F., Munro, E., Banerjee, S. & Gardel, M. L. RhoA Mediates Epithelial Cell Shape Changes via Mechanosensitive Endocytosis. Dev. Cell 52, 152-166.e5 (2020).

21. Berlew, E. E. et al. Single-component optogenetic tools for inducible RhoA GTPase signaling. bioRxiv 2021.02.01.429147 (2021) doi:10.1101/2021.02.01.429147.

22. Yamamoto, K. et al. Optogenetic relaxation of actomyosin contractility uncovers mechanistic roles of cortical tension during cytokinesis. bioRxiv 2021.04.19.440549 (2021) doi:10.1101/2021.04.19.440549.

23. Sasai, Y., Eiraku, M. & Suga, H. In vitro organogenesis in three dimensions: self-organising stem cells. Development 139, 4111–4121 (2012).

24. Lancaster, M. A. & Knoblich, J. Organogenesis in a dish: modeling development and disease using organoid technologies. Science 345, 1247125–1247125 (2014).

25. Clevers, H. Modeling Development and Disease with Organoids. Cell 165, 1586–1597 (2016).

26. Hildebrand, J. D. & Soriano, P. Shroom, a PDZ Domain–Containing Actin-Binding Protein, Is Required for Neural Tube Morphogenesis in Mice. Cell 99, 485–497 (1999).

27. Haigo, S. L., Hildebrand, J. D., Harland, R. M. & Wallingford, J. B. Shroom Induces Apical Constriction and Is Required for Hingepoint Formation during Neural Tube Closure. Curr. Biol. 13, 2125–2137 (2003).

28. Plageman, T. F. et al. Pax6-dependent Shroom3 expression regulates apical constriction during lens placode invagination. Development 137, 405–415 (2010).

29. Chung, M.-I., Nascone-Yoder, N. M., Grover, S. A., Drysdale, T. A. & Wallingford, J. B. Direct activation of Shroom3 transcription by Pitx proteins drives epithelial morphogenesis in the developing gut. Development 137, 1339–1349 (2010).

30. Khalili, H. et al. Developmental Origins for Kidney Disease Due to Shroom3 Deficiency. J. Am. Soc. Nephrol. 27, 2965–2973 (2016).

31. Nishimura, T. & Takeichi, M. Shroom3-mediated recruitment of Rho kinases to the apical cell junctions regulates epithelial and neuroepithelial planar remodeling. Development 135, 1493–1502 (2008).

32. Dietz, M. L., Bernaciak, T. M., Vendetti, F., Kielec, J. M. & Hildebrand, J. D. Differential actin-dependent localization modulates the evolutionarily conserved activity of Shroom family proteins. J. Biol. Chem. 281, 20542–20554 (2006).

33. Hagens, O. et al. A new standard nomenclature for proteins related to Apx and Shroom. BMC Cell Biol. 7, 18 (2006).

34. Hildebrand, J. D. Shroom regulates epithelial cell shape via the apical positioning of an actomyosin network. J. Cell Sci. 118, 5191–5203 (2005).

35. Guntas, G. et al. Engineering an improved light-induced dimer (iLID) for controlling the localization and activity of signaling proteins. Proc. Natl. Acad. Sci. 112, 112–117 (2015).

36. Krueger, D., Tardivo, P., Nguyen, C. & De Renzis, S. Downregulation of basal myosin-II is required for cell shape changes and tissue invagination. EMBO J. 37, e100170 (2018).

37. Araya García, C. A. Formation of Neural Tube. in Reference Module in Biomedical Sciences (Elsevier, 2017). doi:10.1016/B978-0-12-801238-3.11055-4.

38. Massarwa, R., Ray, H. J. & Niswander, L. Morphogenetic movements in the neural plate and neural tube: mouse. WIREs Dev. Biol. 3, 59–68 (2014).

39. Spence, J. R., Lauf, R. & Shroyer, N. F. Vertebrate intestinal endoderm development. Dev. Dyn. 240, 501–520 (2011).

40. Eiraku, M. et al. Self-organizing optic-cup morphogenesis in three-dimensional culture. Nature 472, 51–56 (2011).

41. Okuda, S. et al. Strain-triggered mechanical feedback in self-organizing optic-cup morphogenesis. Sci. Adv. 4, eaau1354 (2018).

42. Lancaster, M. A. et al. Cerebral organoids model human brain development and microcephaly. Nature 501, 373–379 (2013).

43. Gelbart, M. A. et al. Volume conservation principle involved in cell lengthening and nucleus movement during tissue morphogenesis. Proc. Natl. Acad. Sci. 109, 19298–19303 (2012).

44. He, B., Doubrovinski, K., Polyakov, O. & Wieschaus, E. Apical constriction drives tissue-scale hydrodynamic flow to mediate cell elongation. Nature 508, 392–396 (2014).

45. Lee, C., Scherr, H. M. & Wallingford, J. B. Shroom family proteins regulate γ-tubulin distribution and microtubule architecture during epithelial cell shape change. Development 134, 1431–1441 (2007).

46. Campàs, O. A toolbox to explore the mechanics of living embryonic tissues. Semin. Cell Dev. Biol. 55, 119–130 (2016).

47. Ayad, N. M. E., Kaushik, S. & Weaver, V. M. Tissue mechanics, an important regulator of development and disease. Philos. Trans. R. Soc. B Biol. Sci. 374, 20180215 (2019).

48. Ladoux, B. & Mège, R.-M. Mechanobiology of collective cell behaviours. Nat. Rev. Mol. Cell Biol. 18, 743–757 (2017).

49. Hamouda, M. S., Labouesse, C. & Chalut, K. J. Nuclear mechanotransduction in stem cells. Curr. Opin. Cell Biol. 64, 97–104 (2020).

50. Martino, F., Perestrelo, A. R., Vinarský, V., Pagliari, S. & Forte, G. Cellular Mechanotransduction: From Tension to Function. Front. Physiol. 9, (2018).

51. Zimmerman, S. P., Kuhlman, B. & Yumerefendi, H. Engineering and Application of LOV2-based Photoswitches. Methods Enzymol. 580, 169–190 (2016).

52. Tichy, A.-M., Gerrard, E. J., Legrand, J. M. D., Hobbs, R. M. & Janovjak, H. Engineering Strategy and Vector Library for the Rapid Generation of Modular Light-Controlled Protein-Protein Interactions. J. Mol. Biol. 431, 3046–3055 (2019).

53. Woltjen, K. et al. piggyBac transposition reprograms fibroblasts to induced pluripotent stem cells. Nature 458, 766–770 (2009).

54. Lancaster, M. A. et al. Guided self-organization and cortical plate formation in human brain organoids. Nat. Biotechnol. 35, 659–666 (2017).

